# Simple morphometrics for predicting lordosis-induced deviations of body-shape in reared Gilthead seabream (*Sparus aurata* L.)

**DOI:** 10.1101/2021.03.15.435302

**Authors:** Stefanos Fragkoulis, George Koumoundouros

## Abstract

Haemal lordosis, V-shape bending of the haemal vertebrae, is a frequent abnormality of reared fish. Lordosis severity ranges from light deformations of vertebral axis, with insignificant effects on external morphology, to severe axis deformations with significant impact on body-shape. In the present study, we developed a simple morphometric index (PrAn) that links lordosis severity at the juvenile stage with fish body-shape at harvesting, without requiring to radiograph or sacrifice the samples. Examined seabream specimens were part of our previous study (Fragkoulis et al. 2019, Sci. Rep. 9, 9832), which monitored the effects of lordosis on the external morphology of pit-tagged seabream juveniles during their growth, up to harvest size. At both juvenile and adult stages, PrAn was effective in discriminating the normal fish from ca the 70% of lordotic fish. Our results suggest the PrAn as a valuable scale of quality, which quantifies the lordosis effects on fish external morphology, both at the juvenile stage and at harvest. Depending on the lordosis rates, and the hatchery strategy on the maximum allowed abnormality rates, this scale can cull out different rates of lordotic fish, without affecting the fish with normal phenotype or the lordotic fish with high recovery potential.

## Introduction

Skeletal abnormalities develop frequently in reared fish, with a great variability in terms of the affected anatomical area and their effects on fish external morphology. The solution of the problem of skeletal abnormalities relies on a multilevel approach. It involves the identification of abnormalities causative factors, appropriate adjustments of the standard operating procedures in the commercial hatcheries, the standardization of quality control criteria (Koumoundouros 2010, Boglione *et al*. 2013), and the categorization of skeletal abnormalities on the basis of their effects on fish morphology at harvest (Loy *et al*. 1999, de Azevedo *et al*. 2017, Fragkoulis *et al*. 2019).

Haemal lordosis, V-shape bending of the haemal vertebrae, is a frequent abnormality of reared fish (Koumoundouros 2010). It develops during the metamorphosis and early juvenile period, mainly due to elevated swimming activity (Sfakianakis *et al*. 2006a; Palstra *et al*. 2020, Printzi *et al*. 2020). During the juvenile-to-adult growth, lordotic individuals may present a remarkable ability to recover their abnormal phenotype, mainly by repairing the affected vertebrae (Fragkoulis *et al*. 2019). Lordosis severity ranges from light deformations of the vertebral axis, with insignificant effects on external morphology, to severe axis deformations with significant impact on body-shape (Sfakianakis *et al*. 2006b, Fragkoulis *et al*. 2019). Currently, no accurate quality scale exists for the quantification of lordosis severity with respect to its effects on fish phenotype at harvesting. Existing literature is limited in correlating the radiographic appearance of lordosis with juvenile body-shape (Sfakianakis *et al*. 2006b), and in discriminating lordotic juveniles with respect to their recovery potential during the on-growing period by means of a 14-landmark geometric morphometric analysis (Fragkoulis *et al*. 2019).

The goal of the present study was to develop a simple morphometric index that could link lordosis severity at the juvenile stage with fish body-shape at harvesting, without requiring to sacrifice the fish.

## Material and Methods

Examined seabream specimens were part of our previous study (Fragkoulis *et al*. 2019), which monitored the effects of lordosis on the external morphology of pit-tagged (FDX-B, Trovan Ltd, USA) seabream juveniles (154 days post hatching, dph, 86±7 mm standard length, SL) during their growth to harvest size (589 dph, 262±14 mm SL). A total of 146 pit-tagged fish (25 with normal and 121 with lordotic external morphology) were radiographed at the end of on-growing period. As it was shown in Fragkoulis *et al*. (2019), the 54 out of the 146 initially lordotic fish presented a recovered phenotype at harvest. At both sampling ages (154 and 589 dpf), all fish were anaesthetized (ethyleneglycol-monophenylether, Merck, 0.2-0.5 mL L^-1^), photographed on the left side and scanned for ID recognition (Fragkoulis *et al*. 2019).

The effects of haemal lordosis on the body-shape of Gilthead seabream are partially expressed as upward curvature of the caudal peduncle, frequently associated with a shortening of the post-anal body length (Fragkoulis *et al*. 2019). Therefore, we hypothesized that such shape alterations could be detected by an angle (PrAn, predictor angle) defined by the skull (nostril), onset of the post-anal part (anterior base of the anal fin) and the caudal peduncle (base of the central caudal lepidotrichium) (Fig. 1). The XY coordinates of the selected three landmarks were acquired on the digital photographs of the fish by tpsdig2 software (version 2.12, Rohlf 2010) and used in the following formula to estimate the PrAn angle:

**Fig. 1.**
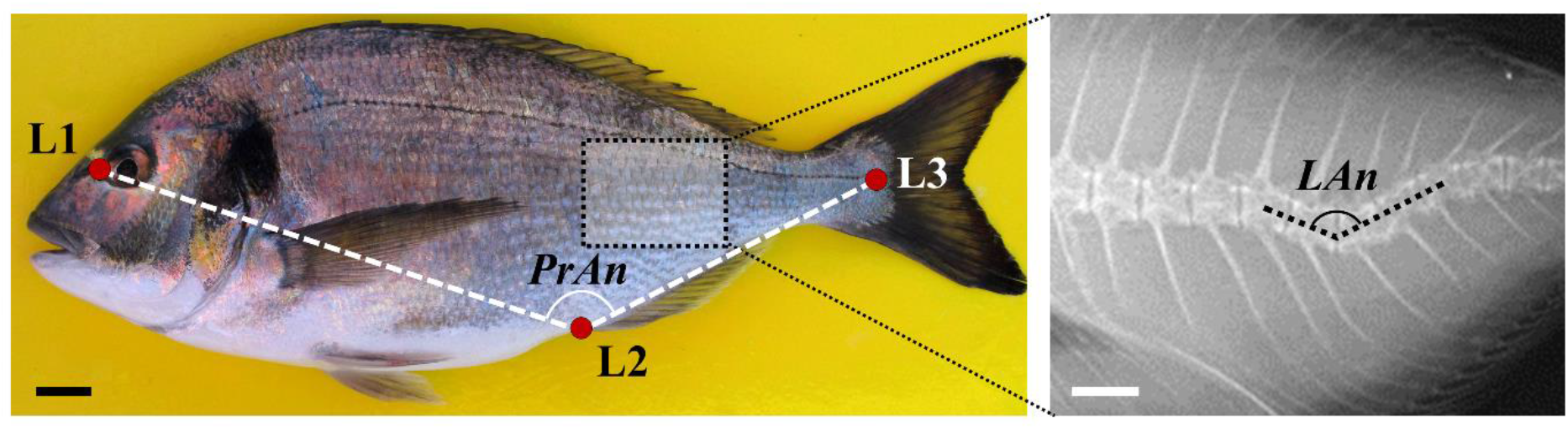
Measurement of the predictor angle (PrAn, left) and the lordosis angle (LAn, right) in adult seabream. The predictor angle was estimated from the X, Y coordinates of three landmark measurements which were positioned on the nostril (L1), the anterior base of the anal fin (L2), and the base of the central caudal lepidotrichium (L3). Lordosis angle was measured on the X-ray of each individual. Scale bars equal to 1 cm.

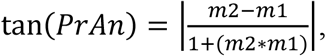

where m2 and m1 is the slope of 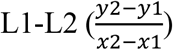 and 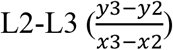 segments respectively (Fig. 1). Moreover, the lordosis angle (LAn) was measured on the x-rays (Fig. 1) of the adult fish (tpsdig2 v.2.12, Rohlf 2010), as a direct estimate of the severity of vertebral bending. The relationship between the predictor angle (PrAn) and the lordosis angle (LAn) was tested by means of piecewise linear regression (Nikolioudakis *et al*. 2010).

## Results

At the adult stage, a significant relationship between the PrAn and LAn was found, with the predictor angle explaining a large amount of the variability in lordosis severity (Fig. 2A). The PrAn-LAn relationship presented a clear inflection point, which separated the normal and recovered fish from the severely lordotic fish (<125.3° PrAn, Fig. 2A). The cumulative frequency distribution of the PrAn revealed that the 70% of the lordotic adults presented PrAn values of <128°, while all the adults with a normal or recovered phenotype had PrAn values of >128 ° (Fig. 2B). At the juvenile stage, PrAn was effective in discriminating the normal fish (Nor, >131°) from the 69% and 72% of lordotic juveniles (<131°) with recovered (Rec) or lordotic (Lor) adult morphology respectively (Fig. 2C). The threshold of 126° PrAn discriminated the 39% of the Lor juveniles from all the Nor and Rec fish (Fig. 2C).

**Fig. 2.**
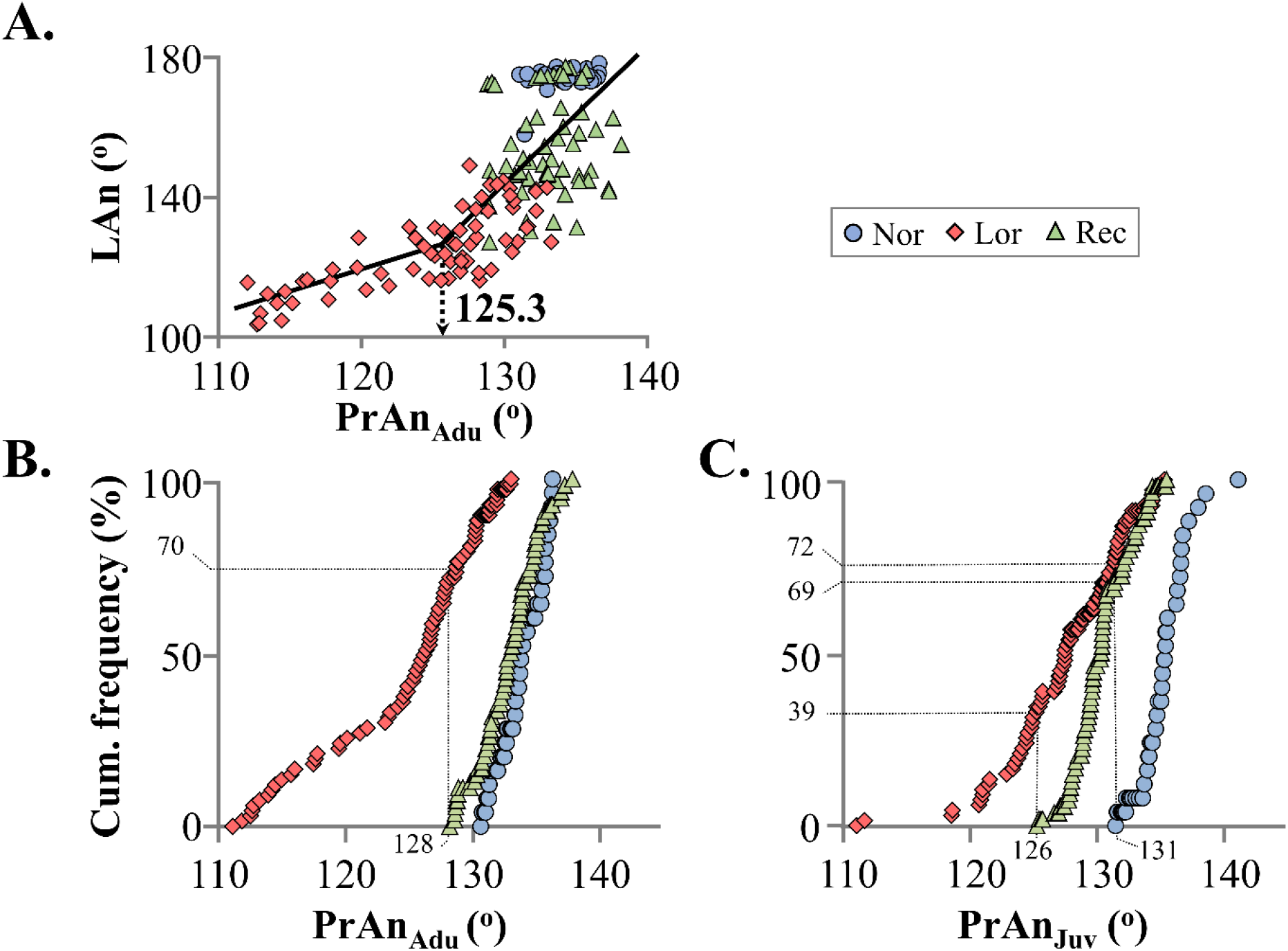
A. Relationship between the lordosis angle (LAn) and predictor angle (PrAn_Adu_) in the adult seabream specimens. **B**. Cumulative frequency of the predictor angle (PrAn _Adu_) for the three phenotypic categories (Nor, Lor, Rec) at the adult stage. **C**. Cumulative frequency of the predictor angle (PrAn_Juv_) for the three phenotypic categories (Nor, Lor, Rec) at the juvenile stage. **Nor**, fish with normal external morphology. **Lor**, fish with lordotic external morphology. **Rec**, fish with a recovery of their lordotic morphology during the juvenile-to-adult growth.

## Discussion

Our results suggest the PrAn as a valuable scale of quality which quantifies the lordosis effects on fish external morphology, both at the juvenile stage and at the harvest. Depending on lordosis rates and the hatchery strategy on the maximum allowed abnormality rates, this scale can cull out different rates of lordotic fish, without affecting the fish with a normal phenotype or the lordotic fish with a high recovery potential (Table S1). Compared with typical geometric morphometric approaches that could be potentially used (Loy *et al*. 1999, Fragkoulis *et al*. 2019), the PrAn index uses fewer landmarks (three) that can be easily measured on anaesthetized fish. The developed quality scale could be incorporated in computerized systems for the automatic quality grading of seabream juveniles, as well as of table-sized fish during the packaging process. In the future, it should be interesting to test the applicability of the developed scale at juvenile smaller than those tested by the present study (e.g. 30-40 mm SL).

## ACKNOWLEDGEMENTS

This study has received funding from the European Union’s Horizon 2020 research and innovation programme under grant agreement No 727610 (PerformFISH). This output reflects only the author’s view and the European Union cannot be held responsible for any use that may be made of the information contained therein.

## DATA AVAILABILITY STATEMENT

The data that support the findings of this study are available from the corresponding authors upon reasonable request.

## REFERENCES

Boglione, C., Gisbert, E., Gavaia, P., Witten, P.E., Moren, M., Fontagne, S. & Koumoundouros, G. (2013). Skeletal anomalies in reared European fish larvae and juveniles. Part 2: main typologies, occurrences and causative factors. Reviews in Aquaculture, 5, S121–S167.

De Azevedo, A.M., Losada, A.P., Barreiro, J.D., Ferreiro, I., Riaza, A., Vázquez, S., & Quiroga, M.I. (2017). Skeletal anomalies in reared Senegalese sole Solea senegalensis juveniles: a radiographic approach, Diseases of Aquatic Organisms, 124, 117–129.

Fragkoulis, S., Printzi, A., Geladakis, G., Katribouzas, N. & Koumoundouros G., (2019). Recovery of haemal lordosis in Gilthead seabream (Sparus aurata L.). Scientific Reports, 9, 9832.

Koumoundouros, G. (2010). Morpho-anatomical abnormalities in Mediterranean marine aquaculture. Transworld Research Network, Kerala, India.

Loy, A., Boglione, C. & Cataudella, S. (1999). Geometric morphometrics and morpho‐anatomy: a combined tool in the study of sea bream (Sparus aurata, sparidae) shape, Journal of Applied Ichthyology, 15, 104–110.

Nikolioudakis, N., Koumoundouros, G., Kiparissis, S. & Somarakis, S. (2010). Defining length-at-metamorphosis in fishes: A multi-character approach. Marine Biology, 157, 991–1001.

Palstra, A.P., Roque, A., Kruijt, L., Jéhannet, P., Pérez-Sánchez, J. & Dirks, R.P. (2020). Physiological Effects of Water Flow Induced Swimming Exercise in Seabream Sparus aurata, Frontiers in Physiology, 11, 1605.

Printzi A., Fragkoulis S., Dimitriadi A., Keklikoglou K., Arvanitidis C., Witten P.E., Koumoundouros G. (2020). Exercise-induced lordosis in zebrafish Danio rerio (Hamilton, 1822). Journal of Fish Biology, 1–8.

Rohlf, F.J. (2010). tpsDig2, Digitize Landmarks and Outlines. In, Department of Ecology and Evolution, State University of New York, Stony Brook, NY, USA.

Sfakianakis, D.G., Georgakopoulou, E., Papadakis, I.E., Divanach, P., Kentouri, A. & Koumoundouros, G. (2006a). Environmental determinants of haemal lordosis in European sea bass, Dicentrarchus labrax (Linnaeus, 1758), Aquaculture, 254, 54–64.

Sfakianakis, D.G., Georgakopoulou, E., Kentouri, M. & Koulmoundouros, G. (2006b). Geometric quantification of lordosis effects on body shape in European sea bass, Dicentrarchus labrax (Linnaeus, 1758). Aquaculture (Amsterdam, Netherlands), 256, 27–33.

