## Supplemental Material for "Simple morphometrics for predicting lordosis-induced deviations of body-shape in reared Gilthead seabream (*Sparus aurata* L.)"

Table S1. Cumulative frequency of fish with lordotic (Lor), recovered (Rec) and normal (Nor) external morphology based on the degrees of the external angle. The Rec group at the juvenile stage consists of the lordotic juveniles that presented recovered external morphology at the adult stage.

| Juveniles | | | | Adults | | | |
|---|---|---|---|---|---|---|---|
| Angle (°) | Lor | Rec | Nor | Angle (°) | Lor | Rec | Nor |
| 112 | **1** | 0 | 0 | 112 | **1** | 0 | 0 |
| 113 | **1** | 0 | 0 | 113 | **6** | 0 | 0 |
| 114 | **1** | 0 | 0 | 114 | **9** | 0 | 0 |
| 115 | **1** | 0 | 0 | 115 | **13** | 0 | 0 |
| 116 | **1** | 0 | 0 | 116 | **16** | 0 | 0 |
| 117 | **1** | 0 | 0 | 117 | **16** | 0 | 0 |
| 118 | **1** | 0 | 0 | 118 | **21** | 0 | 0 |
| 119 | **4** | 0 | 0 | 119 | **21** | 0 | 0 |
| 120 | **4** | 0 | 0 | 120 | **24** | 0 | 0 |
| 121 | **7** | 0 | 0 | 121 | **25** | 0 | 0 |
| 122 | **13** | 0 | 0 | 122 | **28** | 0 | 0 |
| 123 | **15** | 0 | 0 | 123 | **28** | 0 | 0 |
| 124 | **21** | 0 | 0 | 124 | **33** | 0 | 0 |
| 125 | **31** | 0 | 0 | 125 | **40** | 0 | 0 |
| 126 | **39** | 2 | 0 | 126 | **48** | 0 | 0 |
| 127 | 40 | 4 | 0 | 127 | **58** | 0 | 0 |
| 128 | 57 | 11 | 0 | 128 | **70** | 0 | 0 |
| 129 | 60 | 26 | 0 | 129 | 76 | 0 | 11 |
| 130 | 64 | 46 | 0 | 130 | 81 | 0 | 13 |
| 131 | 72 | 69 | 0 | 131 | 90 | 4 | 20 |
| 132 | 84 | 72 | 4 | 132 | 97 | 16 | 35 |
| 133 | 91 | 81 | 8 | 133 | 99 | 28 | 50 |
| 134 | 93 | 91 | 12 | 134 | 100 | 52 | 70 |
| 135 | 97 | 98 | 36 | 135 | | 60 | 81 |
| 136 | 100 | 100 | 60 | 136 | | 88 | 91 |
| 137 | | | 84 | 137 | | 100 | 94 |
| 138 | | | 88 | 138 | | | 100 |
| 139 | | | 96 | 139 | | | |
| 140 | | | 96 | 140 | | | |
| 141 | | | 96 | 141 | | | |
| 142 | | | 100 | 142 | | | |
